# A *Drosophila* model for Dent’s disease reveals impaired ER export of Cubilin as pathogenic mechanism

**DOI:** 10.1101/2025.03.05.641445

**Authors:** Salómon Christer, Zvonimir Marelja, Hannah Hauschild, Marine Berquez, Indira Dibra, Hetvi Gandhi, Yung-Hsin Shih, Martin Helmstädter, Olivier Devuyst, Matias Simons

**Affiliations:** Nephrogenetics unit, Institute of Human Genetics, University Hospital Heidelberg, Heidelberg, Germany; Institut of Physiology, University of Zurich, Zurich, Switzerland; Department of Anatomy and Developmental Biology, Monash University, Melbourne, Australia; Renal Division, Department of Medicine, University Hospital Freiburg, University Faculty of Medicine, Freiburg, Germany

## Abstract

Mutations in the *CLCN5* gene encoding the chloride-hydrogen exchanger ClC-5 cause Dent’s disease, a genetic disorder of the endolysosomal pathway in the proximal tubules of the kidneys. Many patients also develop glomerular lesions, but the underlying mechanism is unclear. We have established an *in vivo* model for Dent’s disease using *Drosophila* nephrocytes that share similarities with podocytes and proximal tubular cells. Upon depletion of *ClC-c*, the fly homologue of *CLCN5*, the endocytic receptor Cubilin was lost from the cortex of nephrocytes, which led to a strong decrease in albumin uptake and slit diaphragm (SD) turnover. Moreover, the actin and microtubular cytoskeleton as well as Rab11-marked recycling endosomes showed a strong cortical accumulation, whereas cholesterol-enriched autophagic compartments emerged in the perinuclear area. Cubilin exhibited a mild mislocalization to cortical early and late endosomal compartments and, in addition, strongly accumulated in the endoplasmic reticulum (ER). This was accompanied by a fragmentation of the ER morphology and an increase in ER exit sites and associated Golgi stacks. These secretory pathway phenotypes were also observed upon silencing of a subunit of the vacuolar H^+^-ATPase (V-ATPase) suggesting that they depend on acidification. Therefore, we speculate that ClC-c and the V-ATPase together acidify the Golgi to allow proper glycosylation and surface trafficking of Cubilin (or its binding partner Amnionless). Interestingly, ER retention of Cubilin was confirmed in ClC-5 knockout mice, underscoring the relevance of this pathomechanism for Dent’s disease.

**Translational statement:** In this work, we study the function of the fly ortholog of *CLCN5* whose mutations cause Dent’s disease, a devastating hereditary kidney disease. By demonstrating that the protein uptake receptor Cubilin is retained in the ER upon ClC-c/ClC-5 depletion in flies and mice, we provide an unexpected new disease mechanism for this disease. Future therapeutic strategies may be directed at improving ER export through acidification of the Golgi apparatus.

## Introduction

Dent disease 1 and 2 (DD1/2) are X-linked recessive disorders of the proximal tubules that are characterized by low-molecular-weight proteinuria, hypercalciuria, nephrocalcinosis and rickets ^1,2^. The main complication of this disease is progression to end-stage renal disease during the second to the fourth decade of life requiring dialysis or transplantation. DD1 is caused by mutations in the chloride channel *CLCN5* and accounts for the majority of cases ^2^, while DD2 is caused by mutations in *OCRL1* ^2,3^. In addition to the tubular phenotypes in DD1, glomerular sclerosis is a quite common histological finding, and more than half of the patients present with nephrotic-range proteinuria ^4,5^. However, the molecular mechanisms underlying the glomerular defects in DD1 have not been explored so far.

ClC-5, the *CLCN5* gene product, is a chloride-proton exchanger localized to the early and recycling endosome (RE) ^6,7^. Cells lacking ClC-5 show decreased levels of the major protein receptors Megalin and Cubilin, leading to tubular proteinuria ^8^. It has been proposed that in ClC-5-deficient cells the receptors are shifted to intracellular compartments, mostly due to impaired endosomal maturation and recycling ^8,9^. In proximal tubule cells, there is a particularly high need for these processes, as the amounts of ligands that need to be taken up and degraded is very high. The acidic pH within early endosomes facilitates the dissociation of internalized ligands from their membrane receptors and is also essential for lysosomal degradation. Acidification is carried out by the vacuolar H^+^-ATPase (V-ATPase), a process that is dependent on chloride-proton exchange by ClC-5. In addition, the chloride ion itself seems to play an undefined role in promoting endocytosis ^10^. Following the dissociation from their bound ligands, membrane receptors are sorted into tubules that can either directly fuse with the plasma membrane via “fast recycling,” or aggregate to form recycling endosomes. As membrane proteins are continuously removed from early endosomes, the remaining spherical structures mature into late endosomes, which eventually deliver their contents to lysosomes. All these events are orchestrated by Rab GTPases, in particular Rab5, Rab7 and Rab11. Rab11, which is responsible for endosomal recycling, also has functions in post-Golgi transport, which are poorly understood ^11–14^. Similarly, the V-ATPase also operates in the Golgi apparatus, which is essential for post-translational modifications of proteins, particularly glycosylation ^15,16^.

In *Drosophila melanogaster*, nephrocytes are renal-like cells that display features of both podocytes and proximal tubular cells ^17,18^. For the filtering and uptake of hemolymph proteins, they are equipped with slit diaphragms (SD) on the cell surface that seal membrane invaginations stretching into the interior of the cell. After filtration into the invaginations, proteins are internalized via the Cubilin/Amnionless (Cubam) tandem receptor (Figure 1A). Under normal conditions, nephrocytes have a concentric organization of the endolysosomal pathway. The most peripheral ring contains early endosomes and invagination membranes, while the second ring harbours the large late endosomes (also called alpha-vacuoles). The lysosomes (beta-vacuoles) can be found in the perinuclear region together with the ER ^19,20^. Finally, REs have been observed throughout the cytoplasm ^21^, but have not yet been studied in detail in nephrocytes. As *ex vivo* nephrocytes preserve the organization and the functionality of the endolysosomal system, the combination of imaging techniques and genetic manipulation allows for a mechanistic dissection of podocyte and proximal tubular diseases. Here, we use *Drosophila* nephrocytes as a model to study ClC-5 function. In addition to several phenotypes within the endolysosomal system, we find that the deficiency of the ClC-5 ortholog, ClC-c, leads to the retention of Cubilin in the ER. As a consequence, Cubilin-dependent endocytosis is shut down, in turn leading to induction of autophagy, reduced SD turnover and a shift of the cytoskeleton towards the cell cortex.

**Figure 1:**
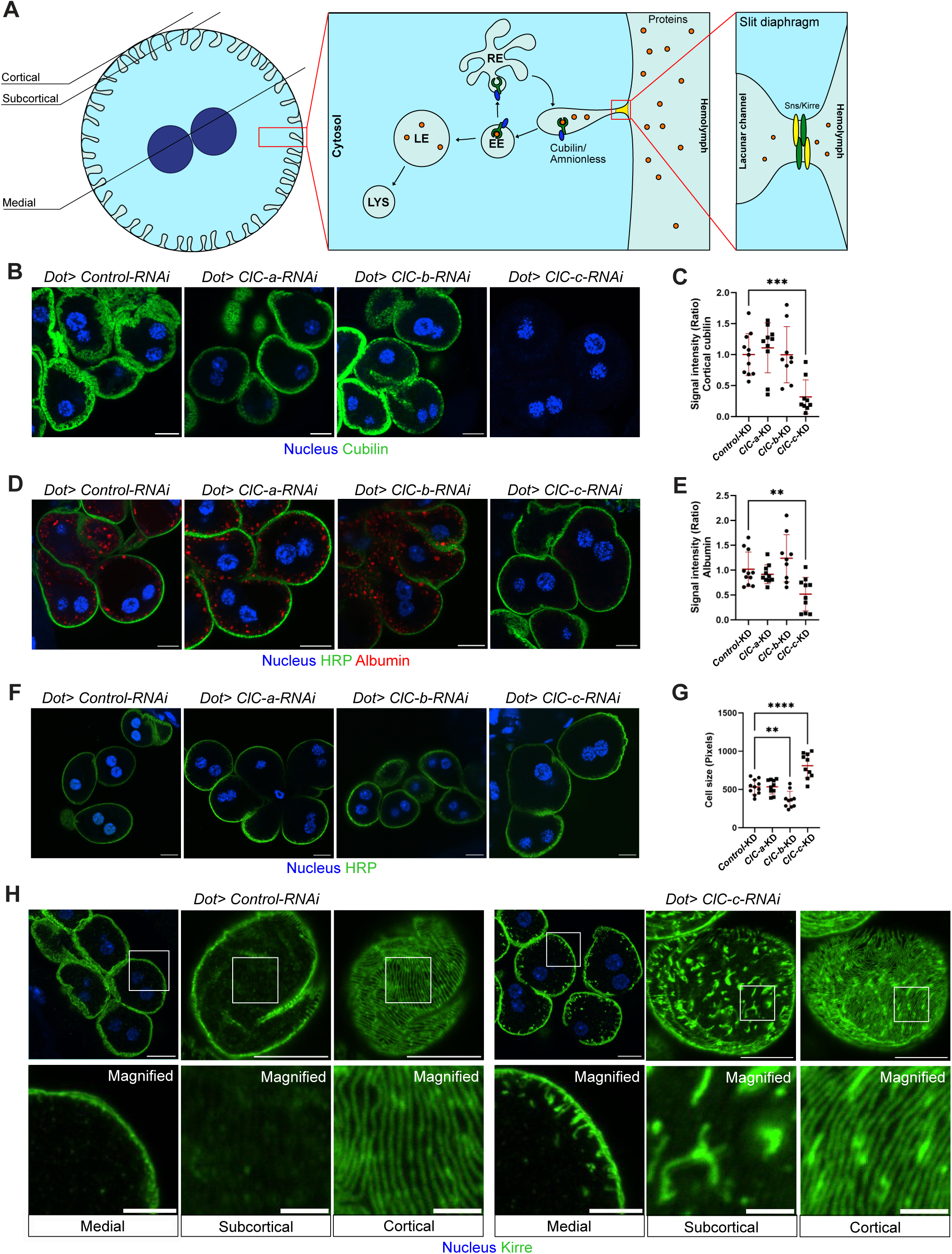
*ClC-c* KD nephrocytes recapitulate Dent’s disease phenotypes. (A) Schematic diagram of *Drosophila* nephrocytes. The cortex of nephrocytes is dotted with membrane invaginations where Cubam (Cubilin/Amnionless) complex is located and mediates endocytosis of proteins filtered by slit diaphragm proteins Sns and Kirre located at the entrance of these invaginations. Following endocytosis, Cubam complex is subsequently transported back to membrane invaginations through the recycling endosomes (RE). Early endosomes (EE) mature into late endosomes (LE) and subsequently lysosomes (LYS) where endocytosed proteins are degraded. (B,C) Non-permeabilized garland nephrocytes expressing RNAi against *ClC-a, ClC-b* and *ClC-c* show loss of cortical Cubilin in upon KD of *ClC-c*. (C) Quantified cortical Cubilin signal mean fluorescence per animal (n ≥ 9) P = 0.0007 for ClC-c-KD. (D,E) Albumin uptake assay with membrane marker HRP performed using the above genotypes shows loss of endocytic activity in *ClC-c* KD nephrocytes. (E) Quantification of albumin signal intensity, Mean fluorescence per animal (n ≥ 9) ANOVA with Dunnet‘s correction: P = 0.0062 for *ClC-c* KD. (F,G) Nephrocytes stained with HRP membrane marker show enlarged nephrocytes upon KD of *ClC-c*, while *ClC-b* KD nephrocytes are mildly diminished in size. Quantification (G) of nephrocyte cell size, average cell size per animal (n ≥ 9) ANOVA with Dunnet‘s correction: P = 0.0052 for *ClC-b* KD, P = <0.0001 for *ClC-c* KD. (H) Control and *ClC-c* KD nephrocytes stained with slit diaphragm (SD) marker Kin of Irre (Kirre) reveal SD localization in the invaginations in *ClC-c* KD nephrocytes. For clarity, the positions of medial, subcortical and cortical sections are depicted in panel A. Medial magnified image scale bar = 5μm, Cortical and subcortical magnified images scale bar = 2μm. Scale bar = 10μm throughout Figures B-F.

## Methods

### Fly strains and Maintenance

*Drosophila melanogaster* crosses were reared in standard food at 25°C. UAS-Gal4 system was used for tissue-specific expression of transgenic-RNAi lines and overexpression. The following stocks were obtained from Bloomington Drosophila Stock Center (BDSC): *Dorothy(dot)-*Gal4 (#6903), UAS-*myr-RFP* (#7118), UASp-RFP.Golgi (#30907) (*GalT-RFP*), UAS-EGFP-myc-2xFyve; UAS-spinster-RFP (#42716), UASp-*GFP-mCherry-Atg8a* (#37749), UAS-*GFP*-RNAi (#9330), UAS-*Cubn-*RNAi (#51736), UAS-*LIMK1*-RNAi (#26294), UAS-*Tor-*RNAi (#33627), UAS-*shi-*RNAi (#28513). The following stocks were used from Vienna Drosophila Resource Center (VDRC): UAS-*ClC-a*-RNAi (110394 KK), UAS-*ClC-b*-RNAi (103420 KK), UAS-*ClC-c*-RNAi (RNAi 1: KK106844) & (RNAi 2: GD6465), UAS-*Rab11-*RNAi (GD22198); UAS-*Rab5-*RNAi (KK103945); UAS-*Rab7-*RNAi (GD40338) UAS-*Vha44-*RNAi (KK101527). *Lamp-3xmCherry* ^22^ and UAS-GFP-Ref(2)P ^23^ were provided by Gabor Juhasz, and *patched(ptc)-*Gal4 and *daughterless(da)-*Gal4 by Norbert Perrimon.

### Immunohistochemistry and microscopy of *Drosophila* tissue

Garland nephrocytes were dissected from third instar larvae in phosphate buffered saline (PBS) and immediately fixed for 20 minutes at room temperature in 4% Paraformaldehyde diluted in PBS (Fixing solution). Wing discs were dissected from pupae 24 hours after pupa formation and fixed as for nephrocytes. Dissected tissues were then permeabilized with three 10-minute washes in PBST (0.3% Triton-X100 in PBS), for non-permeabilized stainings PBS was used instead of PBST. Dissected nephrocytes were subsequently blocked for 20 minutes in blocking solution (4% fetal calf serum in PBST) before incubating overnight at 4°C with 1°-antibodies diluted in blocking solution. The following primary antibodies were used: 1:400 rat anti-Cubilin (Gift from Mar Ruiz-Gómez) ^24^, 1:10 mouse anti-Rab7 (DSHB), 1:300 rabbit anti-Kirre ^25^, 1:1000 rabbit anti-Rab5, 1:5000 rabbit anti-Rbsn5, 1:200 rabbit anti-Rab11 (Gift from A. Nakamura) ^26^, 1:500 mouse anti-PDI (Enzo #ADI-SPA-891-F), 1:500 Rabbit anti-Sec16 (Gift from C. Rabouille) ^27^, 1:500 mouse anti-alpha-Tubulin (Sigma T6199), 1:500 rabbit anti-pS6 (Gift from A.Teleman) ^28^, 1:200 anti-Megalin (Gift from S. Eaton), 1:400 Alexa Fluor™ 555 Phalloidin (Invitrogen, A34055), 1:200 Alexa fluor 488-conjugated goat anti horseradish peroxidase (Jackson Immunoresearch lab #123-545-021). After washing, tissues were incubated for 2 h at room temperature with secondary antibodies (dilution 1:200) in PBST. All nuclei were stained using Hoechst 33342. Tissues were subsequently washed and mounted to slides using Vectashield (Vector laboratories) mounting medium.

Albumin uptake assay performed by dissecting *Drosophila* tissue in Schneider’s medium (S2), pulsed in Tetramethylrhodamine conjugated Albumin (Life Technologies, A23016) solution diluted 0.2mg/mL in S2 medium for 5 minutes in darkness (in wing discs Albumin-488 conjugate (Thermo Fisher A13100) was used instead). All following steps performed in darkness. Sample was subsequently chased 10 minutes in S2 medium, followed by triplicate PBS washes and then subsequently incubated 2 hours with HRP and Hoechst 33342, washed several times in PBS before mounting to slides. LysoTracker™ Red DND-99 assay was performed on nephrocytes and co-stained with Rab7 antibody as previously described ^29^. All confocal microscopy was performed using Nikon AX CSLM microscope using 63x oil objective. Filipin staining was performed by incubating dissected and fixed nephrocytes for 10 minutes in 0.15% glycine in PBS before followed by a 2-hour incubation in Filipin solution (0.05mg/ml Filipin Complex (Sigma, F-9765) in PBS) in darkness. Samples were subsequently washed three times with PBS before mounting to slides, samples were kept in darkness before imaging in Zeiss LSM 780 NLO 2-photon microscope.

For transmission electron microscopy (TEM), nephrocytes were dissected in PBS followed by fixation in a mix of 4% paraformaldehyde, 2% glutaraldehyde, and 0.1 M cacodylate buffer pH 7.4. TEM was carried out using standard techniques ^30^.

### Mouse models

Age-matched mature *Clcn5^Y/+^* (Control) and *Clcn5^Y/-^* mouse littermates (C57BL/6 background) ^31–33^ were maintained under humidity and temperature controlled controlled conditions with 12-hour day/night cycles with free access to standard food. All of the experiments were performed in accordance with the ethical guidelines at University of Zurich and the legislation of animal care and experimentation of Canton Zurich, Switzerland (ZH039/19).

### Immunostaining of mouse kidney sections

Paraffinized kidney sections were prepared according to previously published methods ^34^. Sections were immunostained as previously described ^35^ using following antibodies: 1:1000 rabbit anti-Cubilin (gift from R. Kozyraki) ^36^, 1:100 rat anti-calnexin (Bicell 54036), 1:100 rat anti-Megalin (Bicell Scientific, 31012), mounted to slides and imaged using Nikon AX CSLM microscope using 63x oil objective. Fibrosis was measured using Picro Sirius Red staining kit (Abcam, ab150681) using previously described methods ^37^. A minimum five mice of each genotype were used for every experiment to ensure reproducibility.

### Image analysis

All image analysis was performed using ImageJ software. Mean signal was measured in the medial Z-plane (Nephrocytes showing two nuclei) for consistency. Whole nephrocytes were measured and signal averaged for each animal. For regional measurements of Cubilin signal ROI’s were drawn based on PDI (ER) and HRP (cortex) stainings of garland nephrocyte images. Mean cortical and ER signal intensity of Cubilin staining was then normalized to nuclear Hoechst signal to compensate for potential variance in acquisition. All measurements were averaged for each animal. Co-localization analysis was done via JaCop (BIOP version) plugin in Fiji ImageJ, using thresholding. All experiments were repeated at least 3 times with consistent results. Statistical analysis was performed with 2-tailed, unpaired Student’s t-test for comparison of 2 groups, or one-way ANOVA followed by Tukey’s or Dunn’s multiple comparison test for multiple comparisons after verifying normality. Statistical analyses were performed using the GraphPad Prism 10.0 program. *P* less than 0.05 was considered significant (*p < 0.05, **p < 0.01, ***p < 0.001, ****p < 0.0001). More details on statistics can be found in each figure legend.

### Analysis of mRNA expression in whole larvae

Whole larvae ubiquitously expressing UAS-RNAi with *da*-Gal4 driver were homogenized in lysis buffer and mRNA extracted using RNeasy RNA purification kit (Qiagen 74004) and reverse transcribed using qScript cDNA supermix (Quantabio 95048-025) according to the manufacturer’s instructions. Quality of mRNA assessed using Denovix spectrophotometer. Relative expression was then measured using qPCR which was performed on 96-well PCR plate using (Kit) using qPCRBIO SyGreen® Mix Lo-ROX (PCR biosystems (PB 20.11)) as described by manufacturer and amplified using Quantstudio 3 (Applied Biosystems). Duplicates were performed for each mRNA sample using *alpha-Tubulin* as housekeeping gene. The following primers were used in this publication: *alpha-Tubulin at 84B* (*αTub84B)*: (Fwd: GATCGTGTCCTCGATTACCGC, Rev: GGGAAGTGAATACGTGGGTAGG), *dClC-c* (Fwd: TCCAGTCCAGGCGATGTGAT, Rev: CGCCAGAGTGTCCGAAATCT).

## Results

### ClC-c deficiency in *Drosophila* recapitulates endocytic defects in Dent’s disease 1

The genome of *Drosophila melanogaster* harbours three ClC orthologues, which have been named chloride channel (ClC) -a, -b and -c. As previously shown ^38^, ClC-c shares a high sequence homology to human ClC-5 as well as ClC-3 and -4 (Suppl. Figure 1A,B). To identify which of the three fly orthologues functionally corresponds to human ClC-5 in the kidney, we assessed how silencing of the three individual ClC genes in *Drosophila* garland nephrocytes with *dot-*GAL4 impacts cortical Cubilin localization. Cortical Cubilin localization was measured in non-permeabilized nephrocytes by co-staining a Cubilin-specific antibody with the cortical marker Horseradish peroxidase (HRP) ^39^. While cortical Cubilin levels remained unchanged upon knockdown (KD) of *ClC-a* and *ClC-b*, Cubilin localization was noticeably absent at the cortex of *ClC-c* KD nephrocytes (Figure 1B,C, Suppl. Figure 2A).

To study the consequences of the loss of surface Cubilin, we performed an endocytic uptake assay using fluorescently labelled albumin. *ClC-*c KD nephrocytes showed a significant decrease in intracellular albumin, while albumin levels in *ClC-a* and *ClC-b* KD nephrocytes remained unchanged compared to the control KD (Figure 1D,E). Loss of endocytic uptake was confirmed by a separate RNAi line, and KD efficacy for both RNAi targeting ClC-c was validated via qPCR (Suppl. Figure 2B,C). The reduced uptake of albumin was comparable to what we observed in *Cubilin (Cubn)* KD nephrocytes (Suppl. Figure 2D), suggesting that the albumin uptake defect is likely a consequence of Cubilin loss at the nephrocyte cortex. Additionally, we found that nephrocytes expressing *ClC-c* or *Cubn* RNAi were significantly enlarged (Figure 1F,G; Suppl. Figure 2D), suggesting that the increase of the cell surface might be linked to decreased endocytosis. Given that loss of Cubilin has been shown to affect the endocytic turnover of nephrocyte SDs ^24^, we immunostained *ClC-c*-KD nephrocytes with the SD marker Kin of Irre (Kirre). Similar to the *Cubn* KD (Suppl. Fig 2D), we found that Kirre localized to the nephrocyte surface but also inside the lacunar channels (Figure 1H).

As Megalin is only weakly expressed in nephrocytes ^24,40^, we also tested the effect of silencing *ClC-c* in the pupal wing where Megalin mediates endocytosis of proteins ^40^. Using *ptc-*GAL4 that drives expression at the anterior-posterior boundary of pupal wings, we found that Albumin uptake and Megalin levels were also here suppressed by *ClC-c* KD (Suppl. Figure 2E,F).

Together, these results suggest that ClC-c is the ortholog of human ClC-5 and that *Drosophila* nephrocytes (and pupal wing discs) are suitable models to study both proximal tubular and glomerular phenotypes in DD1.

### Loss of ClC-c leads to ER retention of Cubilin

Previous studies in mice and humans had suggested that Cubilin shows a reduction in surface expression with more intracellular localization in the absence of ClC-5 ^8,41^. Therefore, we performed immunostainings of nephrocytes using Triton X-100 permeabilization. In control cells, Cubilin could not only be detected on the invagination membranes lining the cell surface but also weakly in the perinuclear endoplasmic reticulum (ER), marked by ER marker protein disulfide-isomerase (PDI), which possibly reflects newly synthesized Cubam complexes. Surprisingly, the ER pool of Cubilin was much stronger in *ClC-c* KD nephrocytes (Figure 2A-C). Additionally, the morphology of the ER appeared more fragmented and scattered with occasional juxtaposition of the ER with the cell surface (Figure 2B).

**Figure 2:**
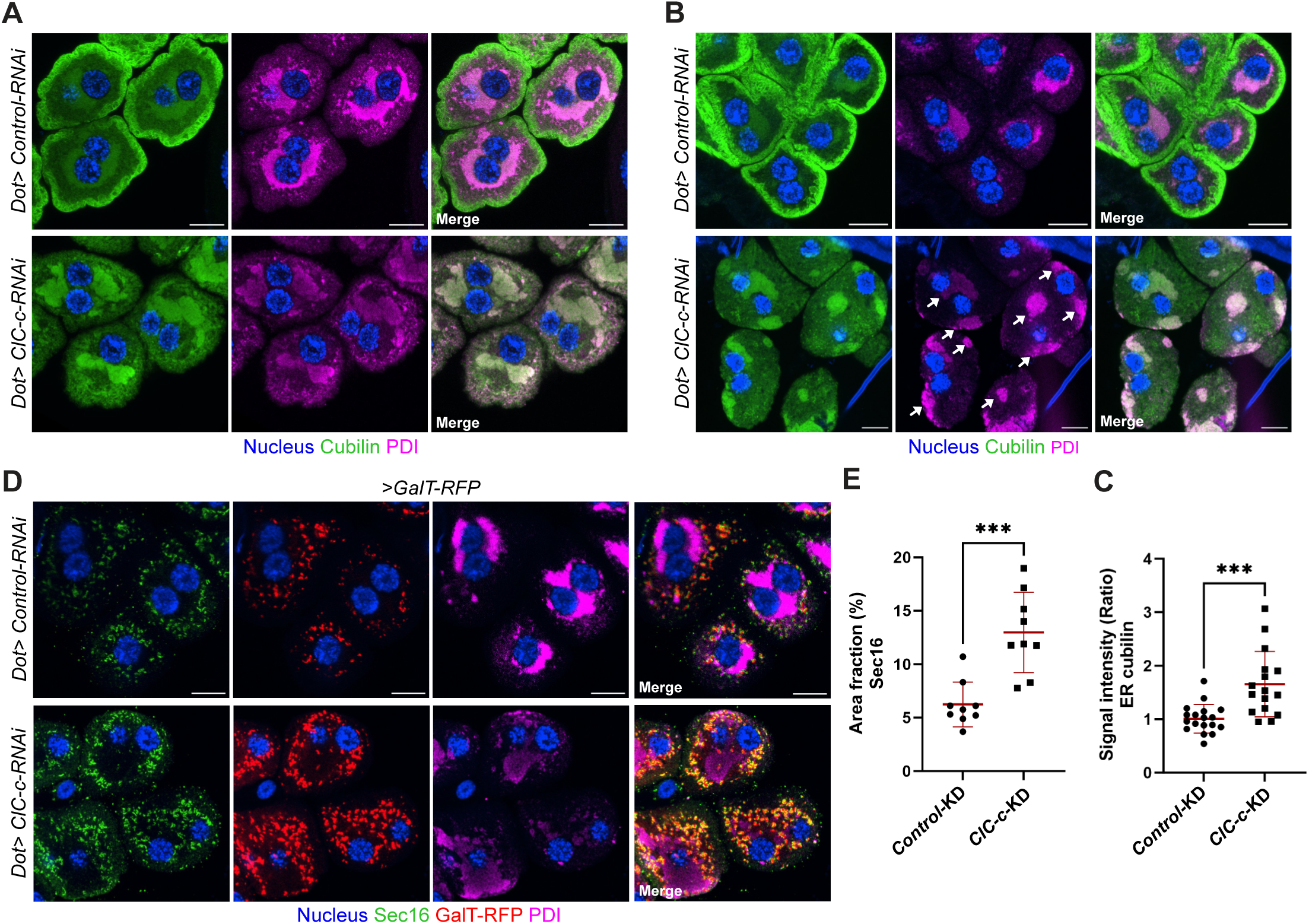
Loss of ClC-c leads to ER retention of Cubilin. Confocal microscopy images of permeabilized control and *ClC-c* KD nephrocytes stained for (A-B) Cubilin and ER marker PDI. (B) Arrows highlight fragmentation of the ER. (C) Quantification of Cubilin signal intensity in area overlapping with PDI shows enhanced Cubilin signal within ER of *ClC-c* KD nephrocytes, averaged signal intensity per animal (n ≥ 20) standard unpaired student’s t-test P = 0.0003. (D,E) *ClC-c* KD nephrocytes expressing RFP-Tagged GalT (trans-Golgi) and stained for Sec16 and PDI show increased ER-exit sites (ERES) and associated Golgi stacks upon loss of ClC-c. (E) Quantified average area fraction of Sec16 (%) per animal (n = 9), Standard unpaired student’s t-test P = 0.0002. Scale bar = 10μm throughout figure.

To further study the early secretory pathway, we stained for the ER-exit site (ERES) marker Sec16 and the trans-Golgi network (TGN) marker Galactosyltransferase-RFP fusion protein (GalT-RFP), which in *Drosophila* cells typically co-localize in the vicinity of the ER. In *ClC-c* KD nephrocytes, the morphology of these ERES-Golgi stacks appeared to be normal compared to control cells but their area coverage was significantly increased (Figure 2D,E). In sum, these observations show that the protein uptake defect in *ClC-c* KD cells is due to ER retention of Cubilin, which in turn is associated with an altered ER morphology as well as an increase in ER exit sites and Golgi stacks.

### ClC-c deficiency causes a strong disorganization of the endolysosomal pathway and the cytoskeleton

We next assessed the impact of *ClC-c* KD on the intracellular organization using transmission electron microscopy (TEM). At the periphery, the displacement of the SDs into the lacunar system could be observed (Figure 3A), reflecting the impaired endocytic turnover of Kirre as shown above (Figure 1H). We also confirmed the fragmentation of the ER and the close apposition of some ER fragments with the plasma membrane leading to the absence of the lacunar channels and the SDs in these areas (Figure 3B). We further observed that alpha-vacuoles/late endosomes were diminished in number and size in the cortical area, while large multilamellar bodies had manifested in the perinuclear region (Figure 3B-D).

**Figure 3:**
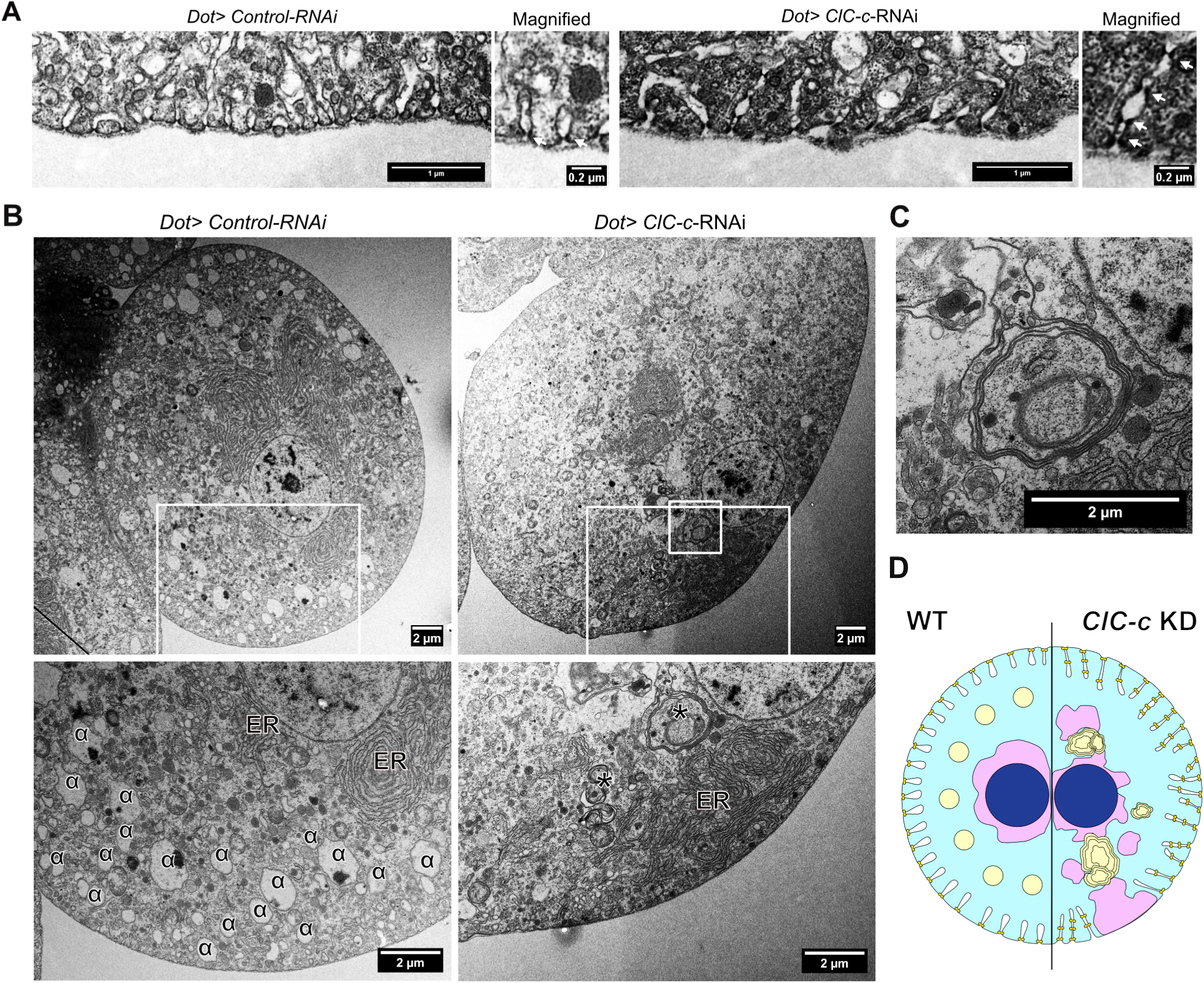
Transmission electron microscopy shows altered surface and intracellular organization in *ClC-c* KD nephrocytes. (A) Transmission electron microscopy (TEM) images show elongated membrane invaginations with ectopic SDs (arrows) in *ClC-c*-KD nephrocytes. Scale bar 1 = μm and 0.2 μm in magnified image. (B) Representative cells with magnifications (lower panels) reveal absence of alpha-vacuoles (α) from cortical area in *ClC-c* KD nephrocytes and formation of multilamellar bodies (labelled with asterisk and magnified in C). Note also perinuclear and cortical localization of ER in control and *ClC-c* KD, respectively. (D) Schematic diagram summarizing the observed phenotypes. Scale bars = 2 μm for figures B-C.

To further study the abnormal endosomal organization, we visualized components of the endo-lysosomal pathway through antibody staining in *ClC-c* KD and control cells. We found that the early endosomal markers Rab5 and Rabenosyn-5 (Rbsn5), which normally occupy the outermost endosomal ring, extended more deeply into the cell interior compared to control cells (Figure 4A, Suppl. Figure 3A). Moreover, the RE marker Rab11 showed a diffuse cytoplasmic localization with some Golgi co-localization in control cells ^11,12,14^, whereas the entire Rab11 pool had shifted to more cortical regions in *ClC-c* KD cells (Figure 4A,C,D). Finally, a very striking effect was found for the late endosomal and lysosomal marker Rab7. While normally marking alpha-vacuoles in the cell cortex, Rab7 re-localized to the perinuclear area upon *ClC-c* KD (Figure 4A). Here, Rab7 decorated large compartments (typically only a few per cell), often between the ER stacks and the nuclei (Figure 4B). Apart from these large compartments, some smaller Rab7-positive vesicles could be observed throughout the cell. While Cubilin signal was noticeably absent from the large perinuclear Rab7 compartment, it did show a partial overlap with the smaller Rab7-positive compartments upon loss of ClC-c, and to an even lesser extent with Rbsn-5 (Figure 4B, Suppl. Figure 3B).

**Figure 4:**
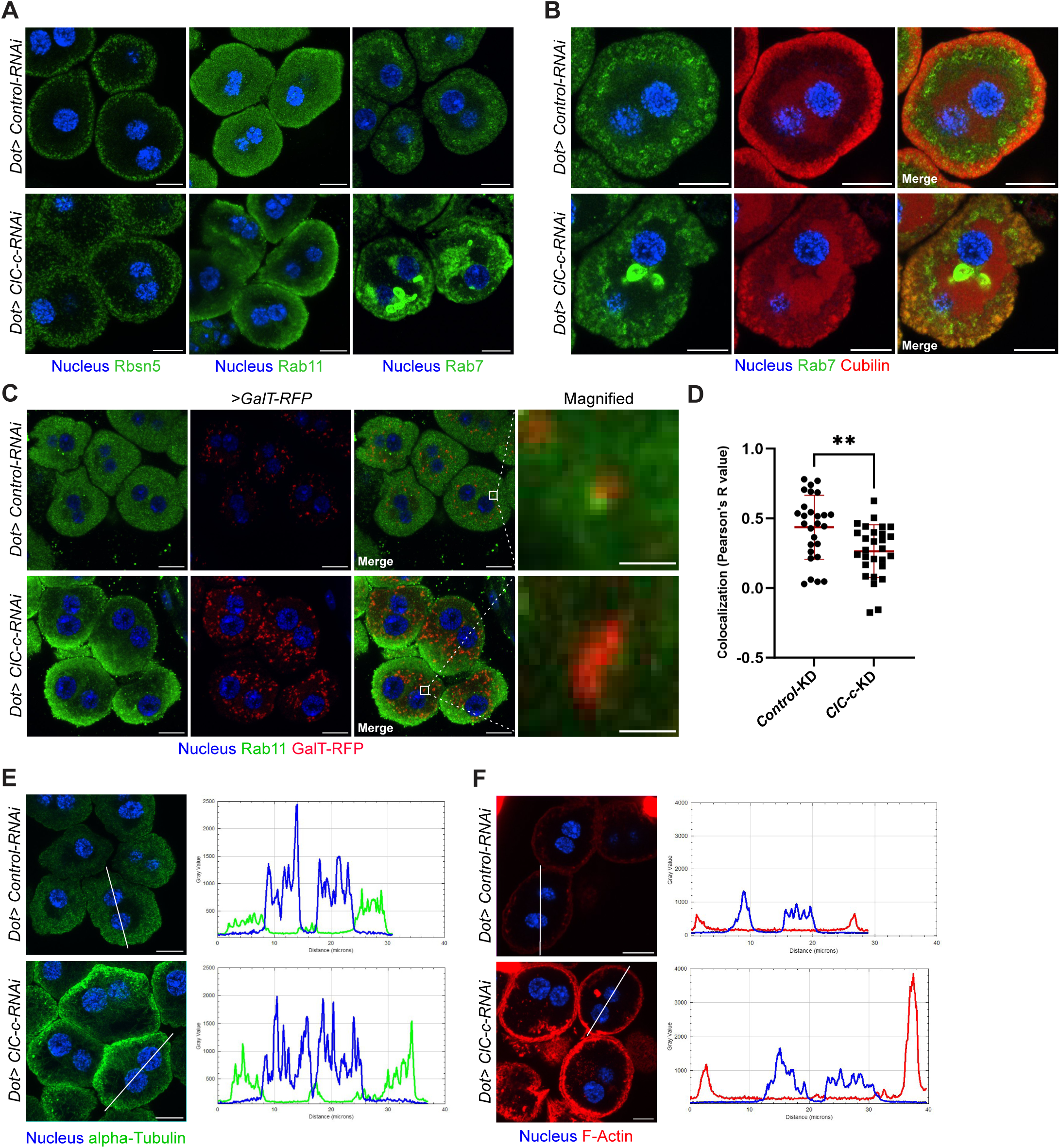
Loss of ClC-c disturbs endolysosomal pathway and leads to cortical build-up of the cytoskeleton. (A-B) Confocal images of control and *ClC-c* KD nephrocytes stained with antibodies for (A) Rbsn5 for early endosomes (EE), Rab11 for recycling endosomes (RE) and Rab7 for late endosomes (LE) and/or lysosomes (LYS). (B) Co-staining for Rab7 with Cubilin show that Cubilin does not co-localize with larger perinuclear Rab7-positive structures, but shows some overlap with smaller Rab7-positive compartments in the cortex. (C,D) Rab11 shift towards cortex in *ClC-c* KD nephrocytes correlates with reduced co-localization of Rab11 with Golgi stacks compared to control. In the magnified inset showing one Golgi stack, the scale bar is 1 μm. In the quantification (D), dots represent individual cells (n ≥ 7) unpaired student’s t-test (P = 0.0037). (E) Antibody staining for alpha-Tubulin reveals enhanced cortical signal in *ClC-c* KD nephrocytes as represented by plot profile. (F) F-Actin (stained by Phalloidin) shows increased cortical signal in *ClC-c* KD nephrocytes, plot profile (Right). Increased diameter of nephrocytes can be attributed to the increased cell size of *ClC-c* KD cells. Scale bar = 10 μm throughout figure.

It has previously been suggested that endocytic impairment in Dent’s disease could be attributed to a disorganised cytoskeletal network ^42–46^. We therefore analysed *ClC-c* KD nephrocytes for any cytoskeletal abnormalities by staining for the actin (Phalloidin) and microtubular (anti-alpha-Tubulin) networks. Both cytoskeletal systems showed a strong shift towards the cortex (Figure 4E,F). Filamentous actin (F-actin) could be observed at the base of the Kirre-marked membrane invaginations (Suppl. Figure 4A) where it aggregated into a barrier-like structure between the plasma membrane and Rbsn-5-positive early endosome (Suppl. Figure 4B). To test whether this actin barrier affected endosomal trafficking, we manipulated the function of twinstar (tsr), which is the ortholog of the actin depolymerization factor Cofilin, a binding partner of ClC-5 ^44^. KD of the negative regulator of tsr, *LIMK1*, resulted in reduced polymerization of actin in *ClC-c* KD nephrocytes (Suppl. Figure 4C,D) but did not have any effect on endocytic uptake indicating that preventing cortical actin accumulation is on its own not sufficient to restore endocytosis (Suppl. Figure 4E,F). These manipulations also did not affect the formation of the Rab7-positive clusters or ER retention of Cubilin (Suppl. Figure 4G-H).

Together, this suggests that ClC-c deficiency leads to an altered organization of the endolysosomal pathway with cortical accumulation of Rab proteins, microtubules and actin. In addition, the absence of ClC-c induced the formation of perinuclear Rab7-positive compartments, which most likely represent the multilamellar structures seen in TEM.

### Cholesterol accumulates in defective autolysosomes of *ClC-c* KD cells

We next sought to further characterize the large Rab7 clusters observed in *ClC-c* KD nephrocytes. Labelling acidic lysosomes using Lysotracker-Red staining revealed a partial overlap with Rab7-positive clusters (Figure 5A). Additionally, overexpression of the lysosomal markers Lamp-mCherry and Spinster-RFP showed similar perinuclear clustering (Suppl. Figure 5A). As the multilamellar structures observed in the TEM suggested lipid accumulation, we performed filipin staining to visualize cholesterol in *ClC-c* KD nephrocytes. This revealed a strong buildup of cholesterol in the Lamp-positive perinuclear clusters (Figure 5B). We also expressed a GFP-mCherry-Atg8a reporter, whose fluorescent tags have different sensitivities for pH ^47^. While GFP-mCherry-Atg8a was generally increased in *ClC-c* KD cells, only the pH-insensitive mCherry was found to be inside the perinuclear compartments, indicating sufficient acidification to quench the GFP signal, in line with the Lysotracker results (Figure 5C). As lysosomal cholesterol accumulation has previously been attributed to defective autophagic flux ^48^, we measured this by expressing a GFP-tagged Ref(2)P (p62 in vertebrates) reporter in *ClC-c* KD nephrocytes. This revealed a strong cytoplasmic accumulation of Ref(2)P in *ClC-c*-KD nephrocytes, indicating impaired autolysosomal clearance (Figure 5D,E) in accordance with previous publications ^49,50^.

**Figure 5:**
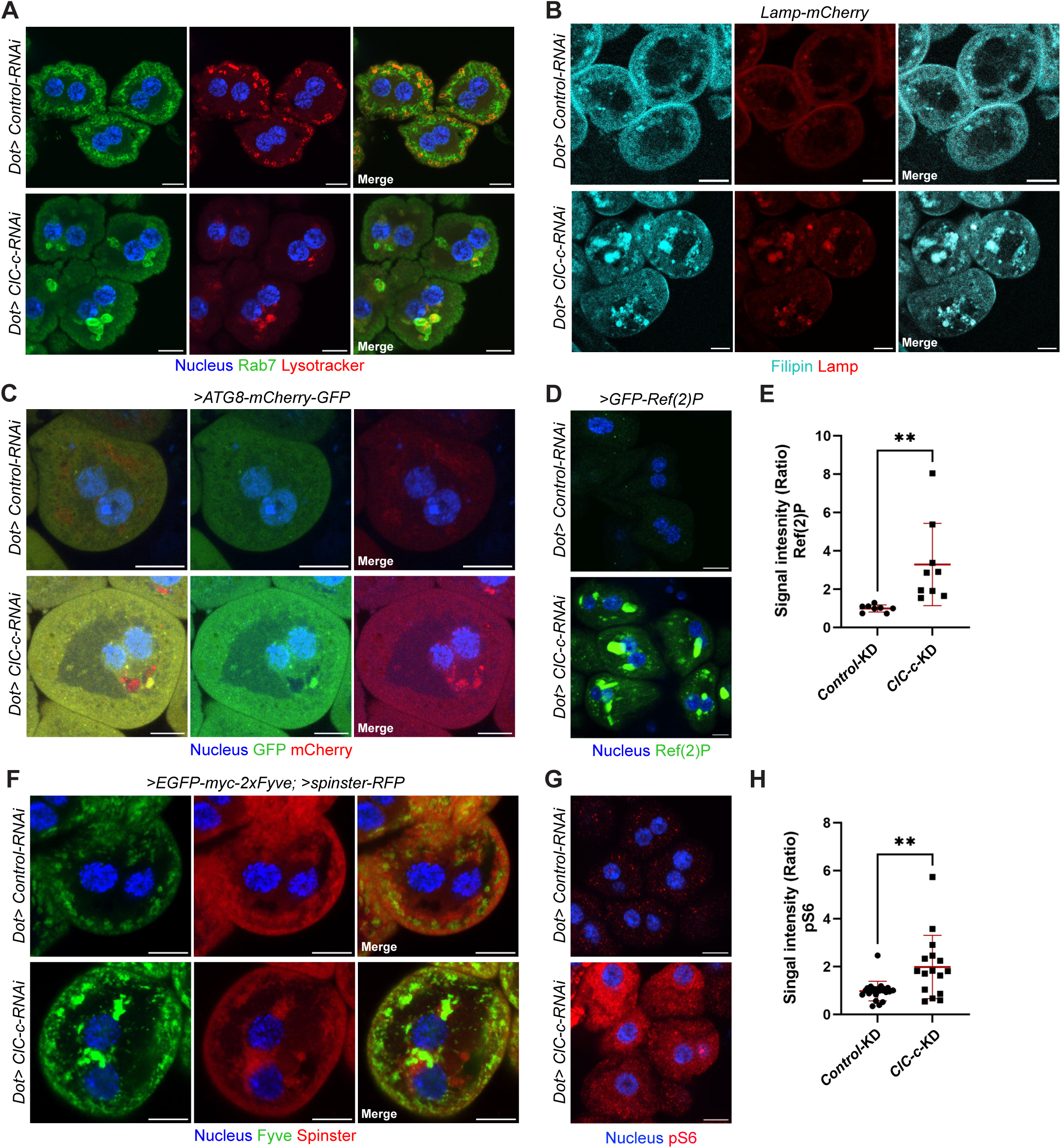
Rab7-positive perinuclear compartments in ClC-c KD cells correspond to autolysosomes. (A) Co-staining of Rab7 and Lysotracker-Red DND-99 (pH<5.5) reveal that perinuclear Rab7-positive clusters in *ClC-c* KD nephrocytes are acidic and correspond to lysosomes. (B) Two-photon confocal microscopy images of Filipin-stained (Cholesterol) nephrocytes co-expressing *ClC-c*-RNAi and lysosomal marker mCherry-tagged Lamp (Lamp-mCherry), shows enhanced cholesterol signal in Lamp-positive clusters. (C) Expression of GFP-mCherry-Atg8a tandem reporter (the GFP-Tag is pH sensitive and thus quenched upon formation of autophagosomes, whereas Atg8a-mCherry signal persists in acidic conditions) reveals formation of acidic autophagosomes in perinuclear region of *ClC-c* KD nephrocytes. (D,E) Overexpression of GFP-Ref(2)P (p62 in mammals) demonstrates impaired degradation and reduced autophagic flux in *ClC-c* KD nephrocytes. Quantification (E) of mean Ref(2)P signal per animal (n ≥ 7) unpaired student’s t-test (P = 0.009). (F*) ClC-c* KD nephrocytes expressing EGFP-myc-2xFyve (to mark PI3P-containing membranes), Spinster-RFP (lysosomal marker) confirm perinuclear clustering of lysosomes as well as re-localization of PI3P-containing membranes to the perinuclear space. (G,H) Activation of mTORC1 upon *ClC-c* KD as measured by antibody staining for phosphorylated-S6 (pS6). Quantification (G) of mean signal intensity of pS6 per animal (n ≥ 7) unpaired student’s t-test (P = 0.0017). Scale bar = 10 μm throughout figure.

To study how autophagy is induced, we visualized the PI(3)P marker eGFP-Fyve that is normally recruited to the perinuclear area to form the phagophore membrane from the ER ^51^. Indeed, PI(3)P-positive membranes emerged at lysosomal perinuclear clusters in addition to its subcortical localization upon loss of ClC-c (Figure 5F). As perinuclear lysosomal positioning and autophagy induction is generally associated with downregulation of mTORC1 ^52^, we also measured the effect of *ClC-c* KD on mTOR activity through antibody staining for phospho-S6 (pS6). Surprisingly, *ClC-c* KD nephrocytes showed significantly increased levels of pS6, corresponding to elevated mTOR activity (Figure 5G,H). However, KD of Tor kinase in *ClC-c* KD nephrocytes, which suppressed S6 phosphorylation, did not alleviate Rab7 clustering or ER retention of Cubilin (Suppl. Figure 5B). This suggests that neither the autophagic nor the ER retention phenotypes are linked to mTOR signalling in *ClC-c* KD cells.

Next, we tested whether decreased endocytosis could lead to autophagy as shown previously for nephrocytes and other tissues ^24,53,54^. For this, we compared the autophagic phenotypes with those caused by the KDs of *Cubn* and *Shibire* (*Shi*; ortholog of dynamin). Indeed, both Cubilin and Shi deficiency exhibited a similar formation of Rab7-positive perinuclear clusters, confirming the relationship between decreased endocytosis and autophagy induction in nephrocytes (Suppl. Fig 6A,B).

Together, these results demonstrate that *ClC-c* KD nephrocytes form large autolysosomal compartments that are unable to clear autophagic cargo, especially cholesterol. The formation of these compartments is mTOR-independent and most likely a result of decreased endocytosis.

### ClC-c deficiency phenocopies Rab11 deficiency in nephrocytes

To better understand what endosomal defect might be responsible for the observed effects on intracellular organization, we compared *ClC-c* KD phenotypes with the KDs of *Rab5, Rab7* and *Rab11*. *Rab5* KD caused a marked loss of cortical Cubilin and accumulation in scattered small punctae. However, these punctae did not co-localize with the ER marker PDI as in the *ClC-c* KD (Figure 6A). KD of *Rab7* led to an accumulation of numerous inflated “empty vacuoles”, as shown previously ^55,56^. These vacuoles fill up the entire cytosolic space leading both to a thin streak of Cubilin at the very apex of the cortex and to a patchy ER with low levels of Cubilin (Figure 6B). By contrast, KD of *Rab11* resulted in loss of Cubilin at the surface and Cubilin retention in a fragmented ER very similar to the *ClC-c* KD (Figure 6C,D). Also here, Rab7 and cholesterol could be found in the large perinuclear clusters (Suppl. Figure 6C-D). Cortical buildup of microtubules (Suppl. Figure C) and an increase in Sec16 punctae could also be observed upon *Rab11* KD (Figure 6E).

**Figure 6:**
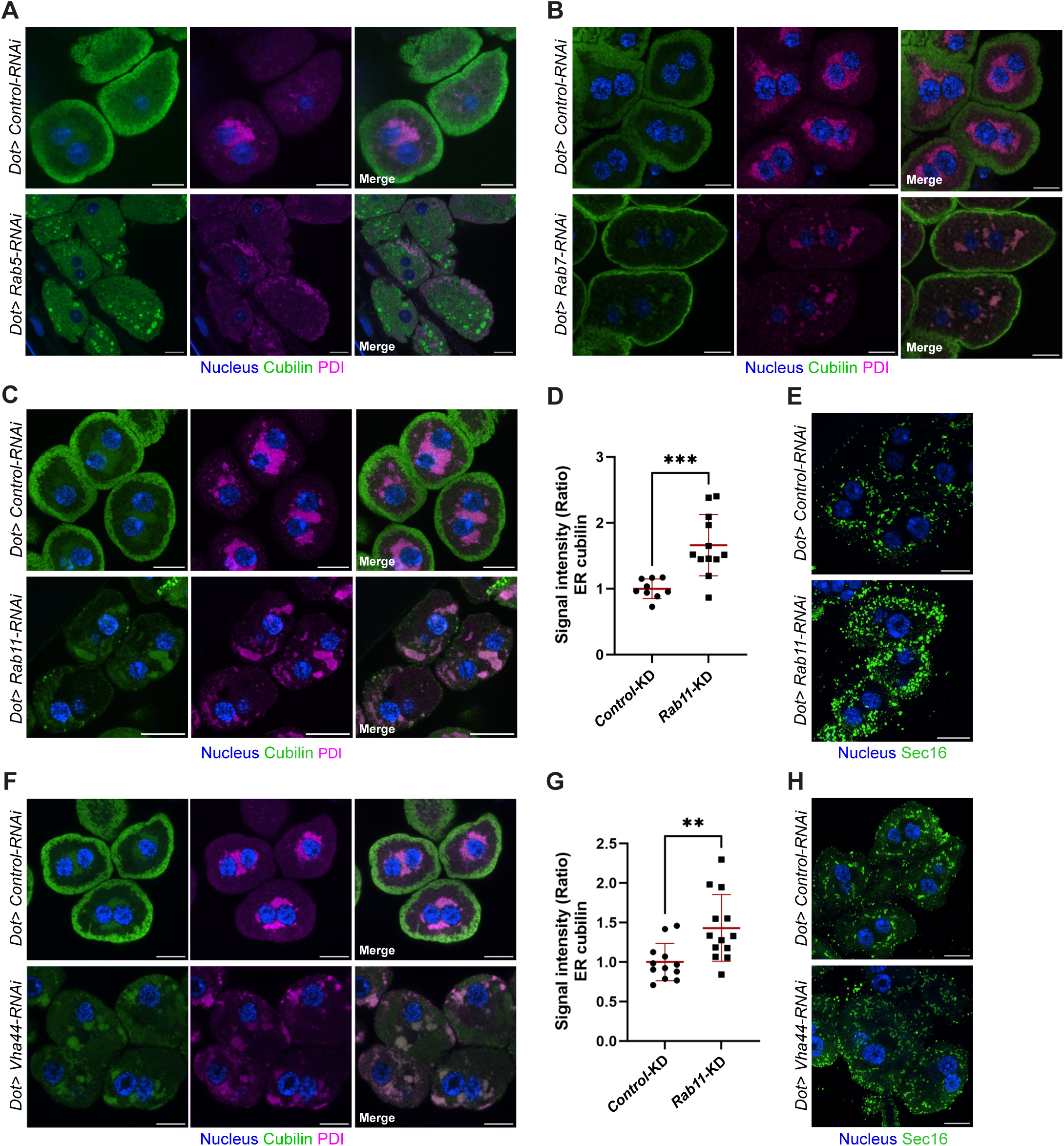
Impaired Rab11 and V-ATPase function phenocopies loss of ClC-c. (A) Confocal images of *Rab5* KD nephrocytes stained with Cubilin and PDI antibodies show strong ER fragmentation and mislocalization of Cubilin to unidentified vesicles. (B) *Rab7* KD nephrocytes show buildup of „empty vacuoles”, more apical localization of Cubilin and disturbed ER morphology. (C,D) *Rab11* KD nephrocytes display retention of Cubilin in a fragmented ER. (D) Quantification of mean ER Cubilin signal intensity per animal (n ≥ 12) unpaired student’s t-test (P = 0.0006). (E) Sec16 antibody staining reveals increased ER-exit sites (ERES) upon loss of Rab11. (F-H) *ClC-c* and *Rab11* KD phenotypes can be recapitulated with depletion of V-ATPase subunit Vha44. (G) Quantification of ER Cubilin signal intensity per animal (n ≥ 12) unpaired student’s t-test (P = 0.0049). Scale bar = 10 μm throughout figure.

Given the role of ClC-5 in acidification, we also silenced V-ATPase subunit *Vha44* (C subunit, ATP6V1C1 in mammals^57^) to measure whether impaired acidification is sufficient to induce the aforementioned phenotypes. Impaired acidification could be validated by diminished Lysotracker seen in *Vha44* RNAi nephrocytes (Suppl. Figure 6D). Antibody staining revealed retention of Cubilin in a fragmented ER and an increase of Sec16 (Figure 6F-H). Moreover, we observed perinuclear Rab7-positive clusters similar to those in *ClC-c* KD cells (Suppl. Figure 6D).

Altogether, the similarities between *ClC-c*, *Rab11* and *Vha44* KD suggest that the functions of these factors are closely connected. As Rab11 and the V-ATPase act both in the Golgi and the endosomal pathway, it can be suspected that ER retention phenotype of Cubilin is caused by a defect in the secretory and not in the endosomal pathway.

### ClC-c phenotypes can be partially reproduced in a mouse model for Dent’s disease

To validate our main Drosophila findings in a mammalian system, we analysed renal cortical sections from *Clcn5^y/-^* mice. These mice have previously been shown to exhibit a number of Dent’s disease features such as low-molecular weight proteinuria, aminoaciduria and glycosuria ^49^. Using Picro-Sirius staining, we observed significant glomerulosclerosis and interstitial fibrosis (Figure 7A,B), which is consistent with findings on renal biopsies of many Dent’s disease patients and a recently generated mouse model with a *CLCN5* mutation ^50^.

**Figure 7:**
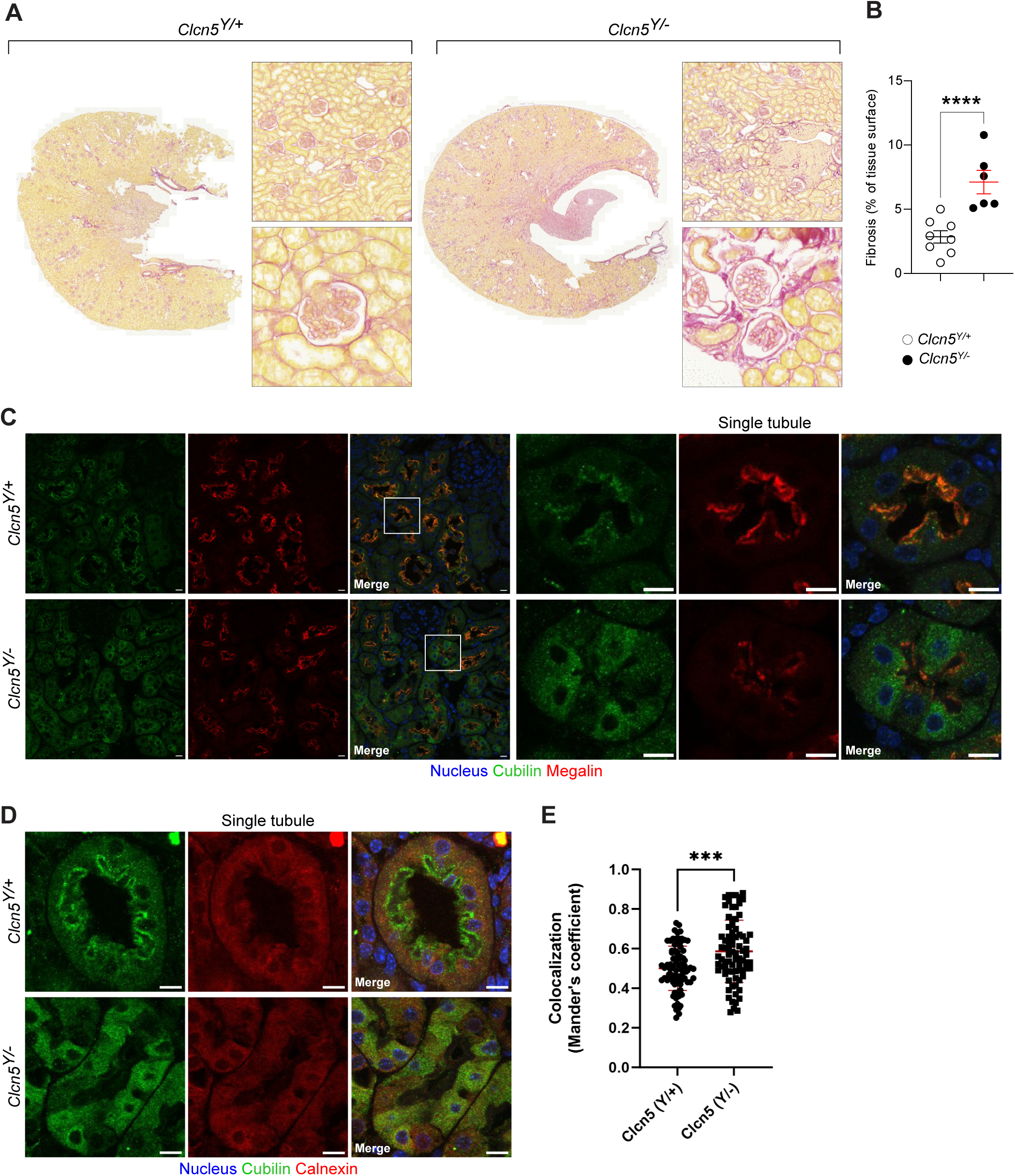
*Clcn5^Y/-^*mice display ER retention of Cubilin in proximal tubules. (A,B) Extracellular matrix deposition measured via Picro sirius red reveals increased fibrosis throughout kidney and enhanced glomerulosclerosis in *Clcn5^Y/-^*compared to *Clcn5^Y/+^* mouse kidney sections. Quantification (B) of mean whole kidney fibrosis per animal (n ≥ 6 kidney sections from mouse of each genotype, (8 *Clcn5^Y/+^* and 6 *Clcn5^Y/-^*mice, total: 14 animals) unpaired student’s t-test (P = 0.0008). (C) Confocal images of paraffinized *Clcn5^Y/+^* and *Clcn5^Y/-^*mouse kidney sections co-stained for Cubilin and Megalin reveals diminished brush border expression of both in the absence of ClC-5. Cubilin also shows enhanced intracellular localization of Cubilin in *Clcn5^Y/-^*mice. (D,E) Co-staining of Cubilin and ER-marker Calnexin reveals overlap between intracellular Cubilin and ER. (E) Quantification of co-localization (Manders coefficient) of Cubilin and Calnexin, each dot represents individual proximal tubule (n = 5 animals, 16 tubule cross sections measured per animal, total 10 animals) unpaired student’s t-test (P = 0.0001).

As reported previously ^8,32^, a significant reduction of brush border Megalin and Cubilin could be observed in proximal tubular cells of *Clcn5^y/-^* mice. However, while Megalin was reduced overall, Cubilin additionally showed increased intracellular localization in *Clcn5^y/-^* proximal tubules (Figure 7C). To determine if the increased intracellularity of Cubilin could be the result of ER retention, we co-stained Cubilin with ER marker Calnexin. This revealed a significant overlap between Cubilin and the ER (Figure 7D,E).

Altogether, our results show that the phenotypes caused by ClC-c deficiency in Drosophila nephrocytes can partially be reproduced in kidneys of mice lacking ClC-5.

## Discussion

Here, we have demonstrated that *Drosophila* is a suitable model to study DD1. While a previous study has studied oxalate crystal formation and ion transport in the fly’s excretory system ^38^, we have focussed on the organellar phenotypes using nephrocytes as a model.

In the past, ClC-5 deficiency was shown to lead to the diversion of the uptake receptors Megalin and Cubilin from the recycling into the lysosomal pathway, where degradation occurs with some delay ^8,9^. This was attributed to impaired early endosomal maturation as a result of defective endosomal acidification ^9^. In the nephrocytes, ClC-c deficiency also led to a partial overlap of Cubilin with early and late endosomes. The main phenotype, however, was the ER retention of Cubilin. In most cells, this phenotype was accompanied by a fragmentation of the ER and an increased number of Sec16-positive ER exit sites and associated Golgi stacks. Given that Rab11 had shifted to cortical regions in the *ClC-c* KD cells, there was also significantly less Rab11 in the vicinity of these structures, which may explain the phenocopy by *Rab11* KD. Rab11 has been shown to regulate post-Golgi transport in addition to its role in endosomal recycling ^11–14^. Therefore, it is possible that the Cubilin retention phenotype in *ClC-c* KD cells might stem from a defective secretory and not from a defective endosomal recycling pathway. Another reason for the ER retention could be the cortical accumulation of the cytoskeleton. In particular the microtubules, which also in the *Rab11* KD showed a cortical accumulation, may be necessary for the transport of Cubilin-containing vesicles from the ER to the surface ^58^.

The phenocopy achieved with KD of *Vha44* might also suggest a problem with the acidification of the Golgi. Defective Golgi acidification due to mutations in V-ATPase subunits has been associated with human glycosylation disorders, causing defects in protein secretion ^16^. As N-glycosylation is critical for cell surface targeting of Cubilin and Amnionless ^59^, defective Golgi acidification and subsequent under- or misglycosylation of the Cubam complex might contribute to the ER retention phenotype. This effect may also be secondary due to mistrafficking of glycosyltransferases as shown for Rab11 silencing ^60^. Given that surface transport of the transmembrane protein Kirre was still possible in *ClC-c* KD cells, it could, however, be that the Cubam complex has different requirements for surface transport than other proteins.

Another striking phenotype in *ClC-c* KD nephrocytes was the strong induction of autophagy with formation of large autolysosomes in the perinuclear area. The similarity of the autophagic phenotype with those seen in *Cubn* and *Shi* KD cells points toward defective endocytosis as a stimulus for autophagy. By contrast, other autophagy-inducing conditions, such as mTOR inhibition and cytoplasmic protein aggregation, led to quite different autophagic phenotypes in nephrocytes ^61,62^. Interestingly, as Cubilin was localized to the surface in *Shi* KD cells and Cubilin was never found in the autolysosomes in *ClC-c* KD cells, there does not seem to be a relation between autophagy and the ER retention of Cubilin. Although previous results have suggested reduced lysosomal acidification in *CLCN5* KO cells ^9^, we were able to observe both a Lysotracker signal and an mCherry signal from the Atg8a tandem reporter inside the large autolysosomal compartments. But as we did not measure absolute pH levels, we cannot rule out that maximal acidification is disturbed. Still, the accumulation of p62 argues for a disturbed autophagic clearance in *ClC-c* KD cells, which may also be caused by the accumulation of cholesterol.

In recent years, it has become clear that DD1 not only affects proximal tubules. Several reports show that many Dent’s disease patients can have nephrotic-range proteinuria. Histology data from renal biopsies as well as ClC-5 expression studies strongly suggest a podocyte involvement ^5,63,64^. In this regard, it is interesting that we observed SD defects upon ClC-c deficiency. Overall, they were very similar to the previously reported SD phenotypes related to Cubilin deficiency in nephrocytes ^24^. In this case, the ectopic SDs in the invaginations were explained by a decreased endocytic turnover in the presence of unchanged exocytic trafficking of SD proteins to the invagination membranes. Thus, it seems that the downregulation of Cubilin upon *ClC-c* KD is not only responsible for reduced albumin uptake but also for reduced SD endocytosis. Whether Cubilin-dependent SD turnover also occurs in mammalian podocytes remains to be determined. However, given that patients with *CUBN* mutations have much milder kidney phenotypes than Dent’s disease patients ^65^, this can at least be questioned.

Overall, our study provides novel insights into the pathogenesis of Dent’s disease. According to our results, Dent’s disease is a disease with multiple cellular phenotypes. Our most unexpected finding is the ER retention of Cubilin that we could confirm in *Clcn5* KO mice. Based on the similarity with the *Vha44* and *Rab11* KD, we speculate that this phenotype is connected with impaired Golgi function. While the yeast chloride-proton exchanger Gef1p has been shown to functionally interact with the V-ATPase for Golgi acidification ^66^, only ClC-3B has so far been shown to localize to the Golgi in mammals ^67^. Therefore, a more detailed analysis of the Golgi pH in the absence of ClC-c and ClC-5 will be necessary in the future. A better understanding of the mechanisms underlying Cubilin’s ER retention may therefore provide general insights into the role of ClC exchangers, proton pumps and Rab11 along the secretory pathway between species.

## Disclosure statement

All the authors declared no competing interests.

## Data sharing statement

All data supporting the findings used in this study are present in the main figures and supplementary material. Additional data may be requested from corresponding author upon reasonable request

## Acknowledgements

We are grateful to Svenja Keller for technical help and to Severine Kayser for excellent technical assistance with TEM. We thank Mar Ruiz Gomez, Akira Nakamura, Catherine Rabouille, Aurelio Teleman and Suzanne Eaton for providing antibodies as well as Norbert Perrimon, Gabor Juhasz, the Bloomington stock center and Vienna Drosophila RNAi Center (VDRC) for providing flies.

## Funding Statement

We acknowledge the European Research Council (ERC) under the European Horizon 2020 research and innovation programme (Grant agreement No. 865408 (RENOPROTECT), the Deutsche Forschungsgemeinschaft (DFG SI1303/5-1 (Heisenberg-Programm) and DFG SI1303/6-1), the Steno Collaborative Grant from the NovoNordisk Foundation (NNF18OC0052457) (all to M.S.). The Core Facility for Electron Microscopy (EMcore) at the University Freiburg Medical Center—IMITATE is registered with the DFG (German Research Foundation) under the reference number RI_00555. OD is supported by the Swiss National Science Foundation (grant 320030-232229), and the University Research Priority Program (URPP) ITINERARE at the University of Zurich.

## Author Contributions

SC, ZM, HH, MB, YS, ID, MH performed experiments and analyzed data. OD, MS provided reagents, performed data analysis. MS conceived the study and wrote the paper together with SC. All authors performed manuscript editing.

## Notes

### Competing Interest Statement

The authors have declared no competing interest.

