## Supplementary Figures for "A *Drosophila* model for Dent’s disease reveals impaired ER export of Cubilin as pathogenic mechanism"

**Suppl. Fig 1:**

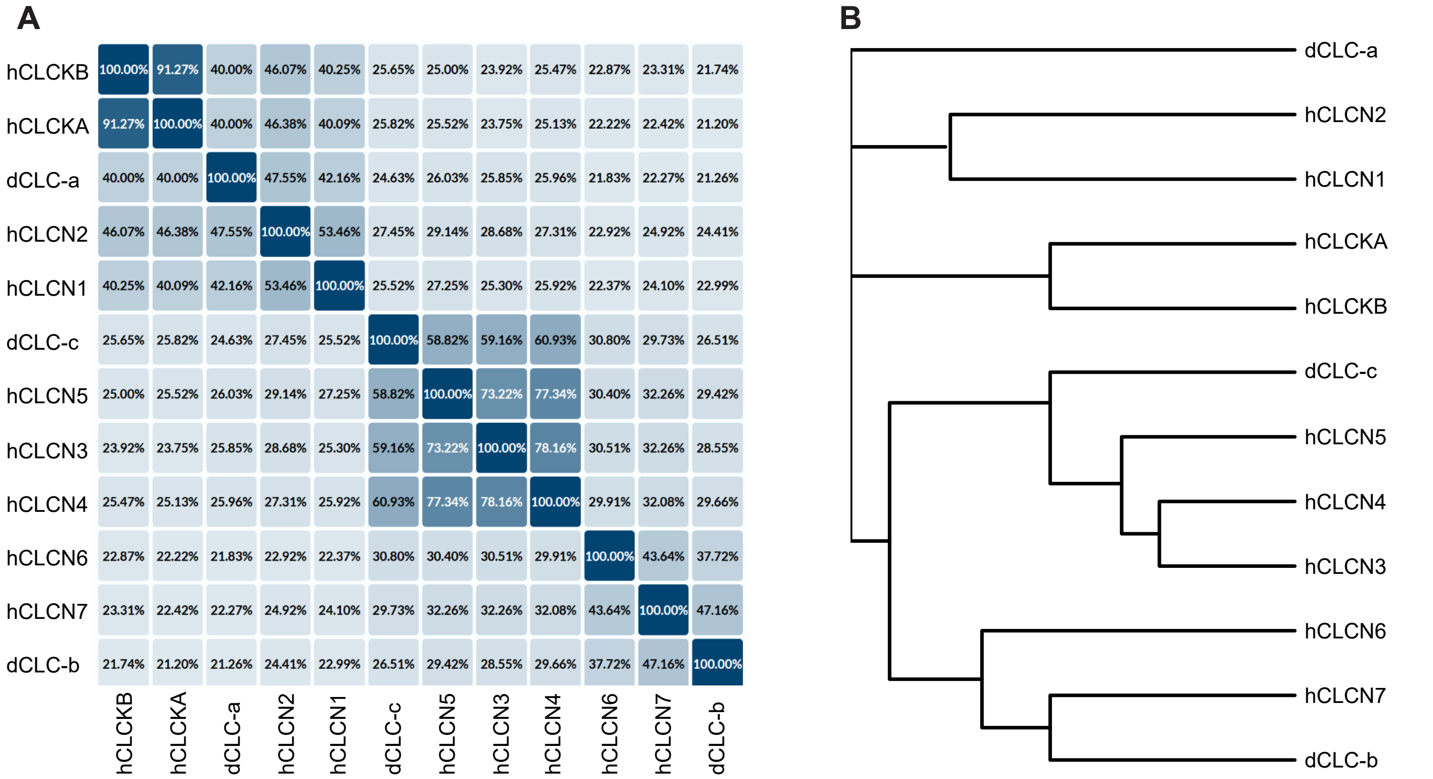


**Suppl. Figure 1: Amino acid sequence alignment between *Drosophila* ClC-c and human ClC-5.** (A) Homology chart displaying percentage of amino acid homology between human and *Drosophila* ClC family proteins. (B) Phylogenetic tree of ClC family proteins. Homology chart and phylogenetic tree generated using Uniprot protein alignment tool and adapted in Affinity Designer.

**Suppl. Fig 2:**

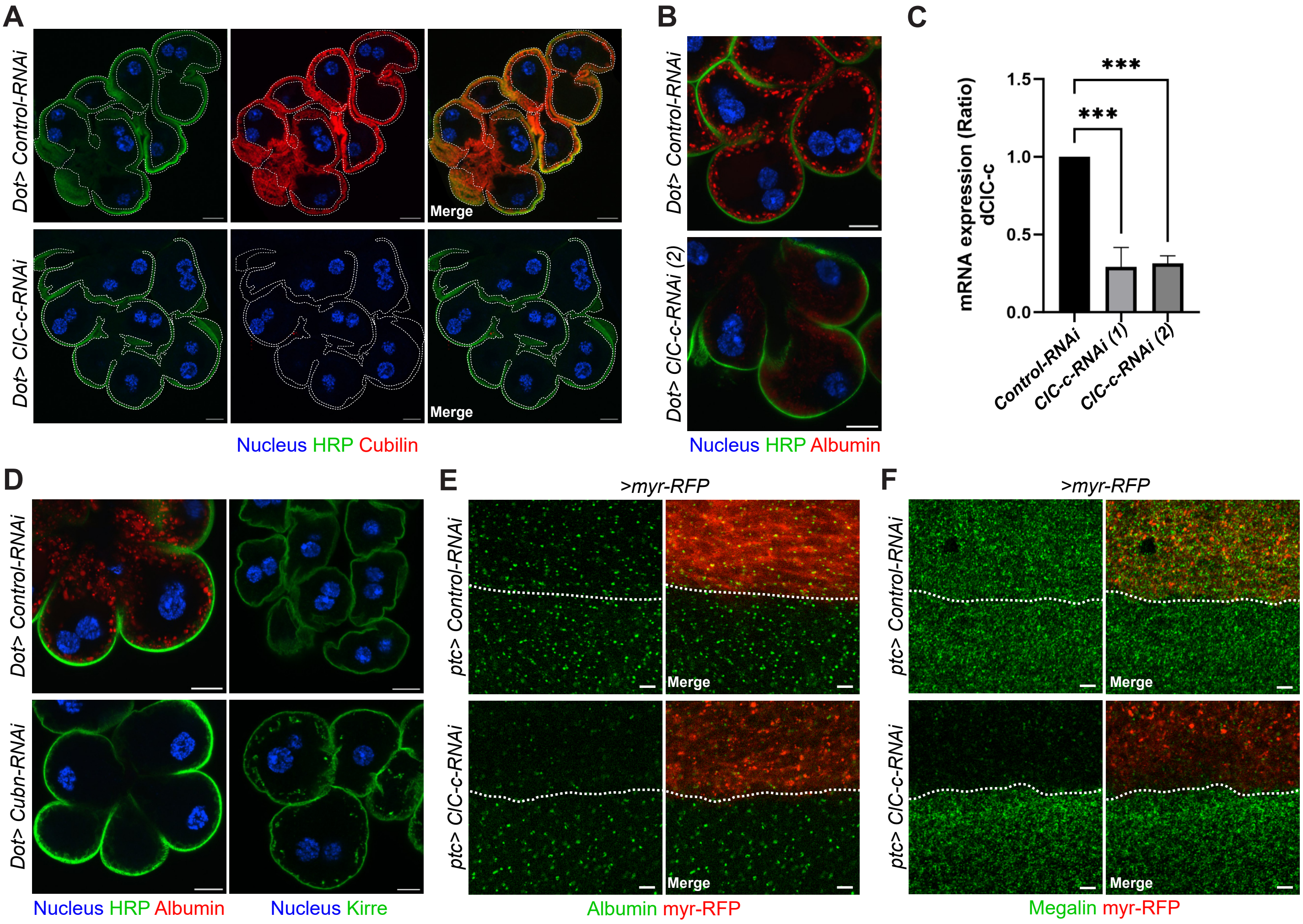


**Suppl. Figure 2: *ClC-c* and *Cubn* silencing in nephrocytes and pupal wings.** (A) Confocal microscopy images of non-permeabilized *ClC-c* KD garland nephrocytes stained for Cubilin and cortical marker horseradish peroxidase (HRP). Dotted lines indicate selected cortical area (based on HRP signal) used to quantify cortical Cubilin localization in Figure 1B. (B) Albumin uptake using an alternative ClC-c RNAi construct (*ClC-c*-RNAi (2)). C) Both *ClC-c* RNAi contructs (1 & 2) validated via RT-qPCR of mRNA from whole larvae expressing RNAi constructs with *daughterless*-GAL4. Relative expression ratio calculated using the ΔΔCt method, (n = 3) unpaired ANOVA with Dunnet's correction P = 0.0010 for both RNAi constructs. (D) Loss of endocytic uptake in *Cubn* KD nephrocytes via Albumin uptake assay + HRP staining and reduced slit diaphragm turnover visualized by antibody staining for Kirre (E-F) Confocal microscopy images of *patched-Gal4* driven co-expression of RFP and *Control* or *ClC-c* RNAi in the anterior posterior boundary of pupal wings. (E) *ClC-c* KD cells (RFP-positive) show reduced endocytic uptake of Albumin. (F) Staining with Megalin-specific antibody reveals reduced expression of Megalin in *ClC-c*-KD pupal wing epithelium. Scale bar = 10µm throughout figure.

**Suppl. Fig 3:**

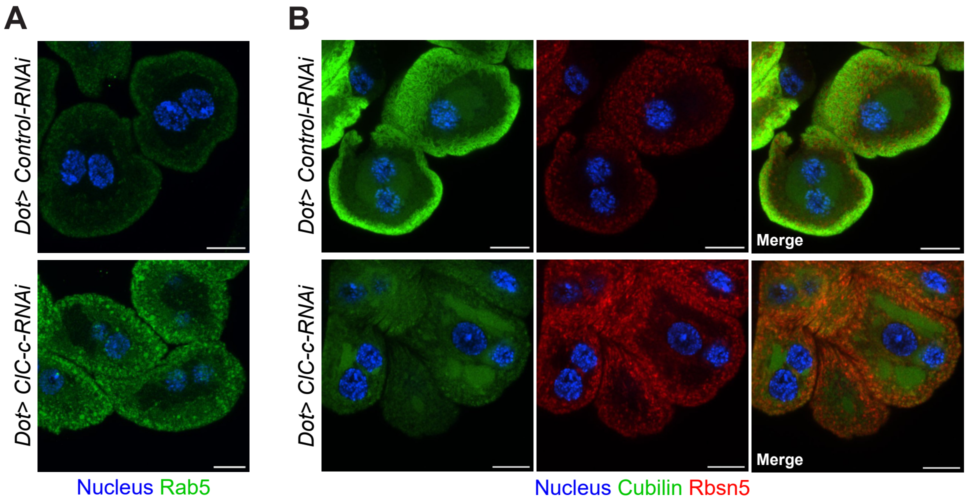


**Suppl. Figure 3: Co-staining of Cubilin and early endosomes in *ClC*-c KD nephrocytes.** Confocal microscopy images of dissected garland nephrocytes (A) stained with Rab5 antibody reveals mislocalization of early endosomes (EE). Co-staining of Rbsn5 (EE) and Cubilin shows some overlap between Cubilin and EE in *ClC-c*-KD nephrocytes. Scale bar = 10µm throughout figure.

**Suppl. Fig 4:**

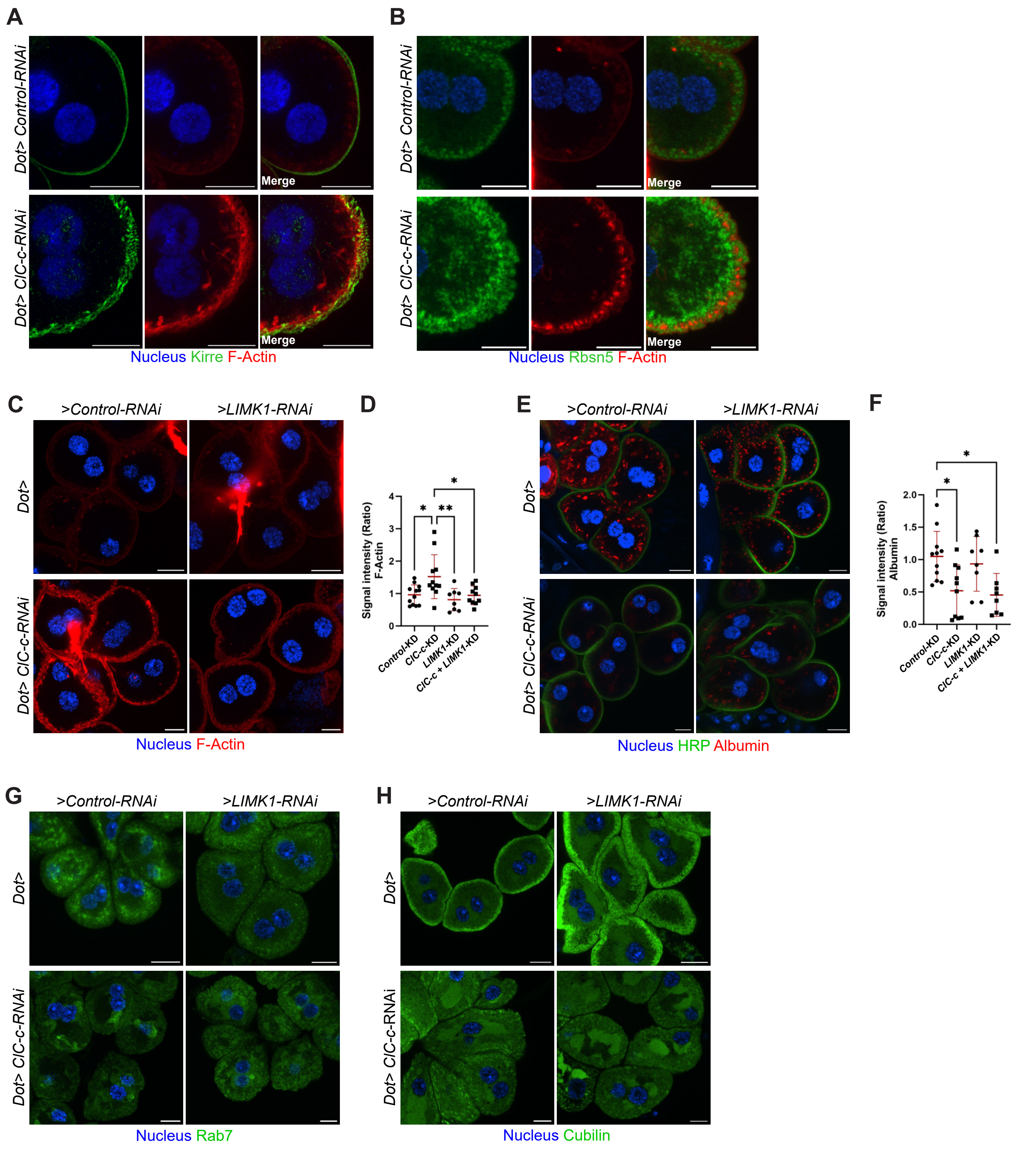


**Suppl. Figure 4: Cortical actin accumulation does not contribute to Dent's disease phenotypes in *ClC-c* KD nephrocytes.** (A) Co-staining for Kirre and F-actin (Phalloidin) revealed actin build-up at the base of the invaginations in *ClC-c*-KD nephrocytes (B) Co-staining for F-Actin and Rbsn5 (early endosomes (EE)) reveals that subcortical actin is localized within broadened EE region (C) Silencing of *LIMK1* rescues cortical F-actin accumulation in *ClC-c* KD nephrocytes. (D) Quantification of mean F-actin signal per animal ($n \geq 8$), unpaired ANOVA with Tukey's correction ($P = 0.0211$ Control vs *ClC-c* KD, $P = 0.0070$ *ClC-c*-KD vs *LIMK1*-KD, $P = 0.0240$ *ClC-c*-KD vs *ClC-c*-KD + *LIMK1*-KD). (E) Albumin uptake assay performed on the same genotypes with cortical marker HRP reveals unaltered uptake of Albumin upon KD of *LIMK11* in *ClC-c* KD cells. (F) Quantification of mean Albumin signal per animal ($n \geq 8$), unpaired ANOVA with Tukey's correction ($P = 0.0183$, Control vs *ClC-c*-KD; $P = 0.0119$, *Control* vs *ClC-c*-KD + *LIMK1*-KD). Nephrocytes co-expressing *ClC-c* and *LIMK1* RNAi show no change in Rab7 (G) and Cubilin (H) localization compared to *ClC-c* KD alone. Scale bar = 10µm

**Suppl. Fig 5:**

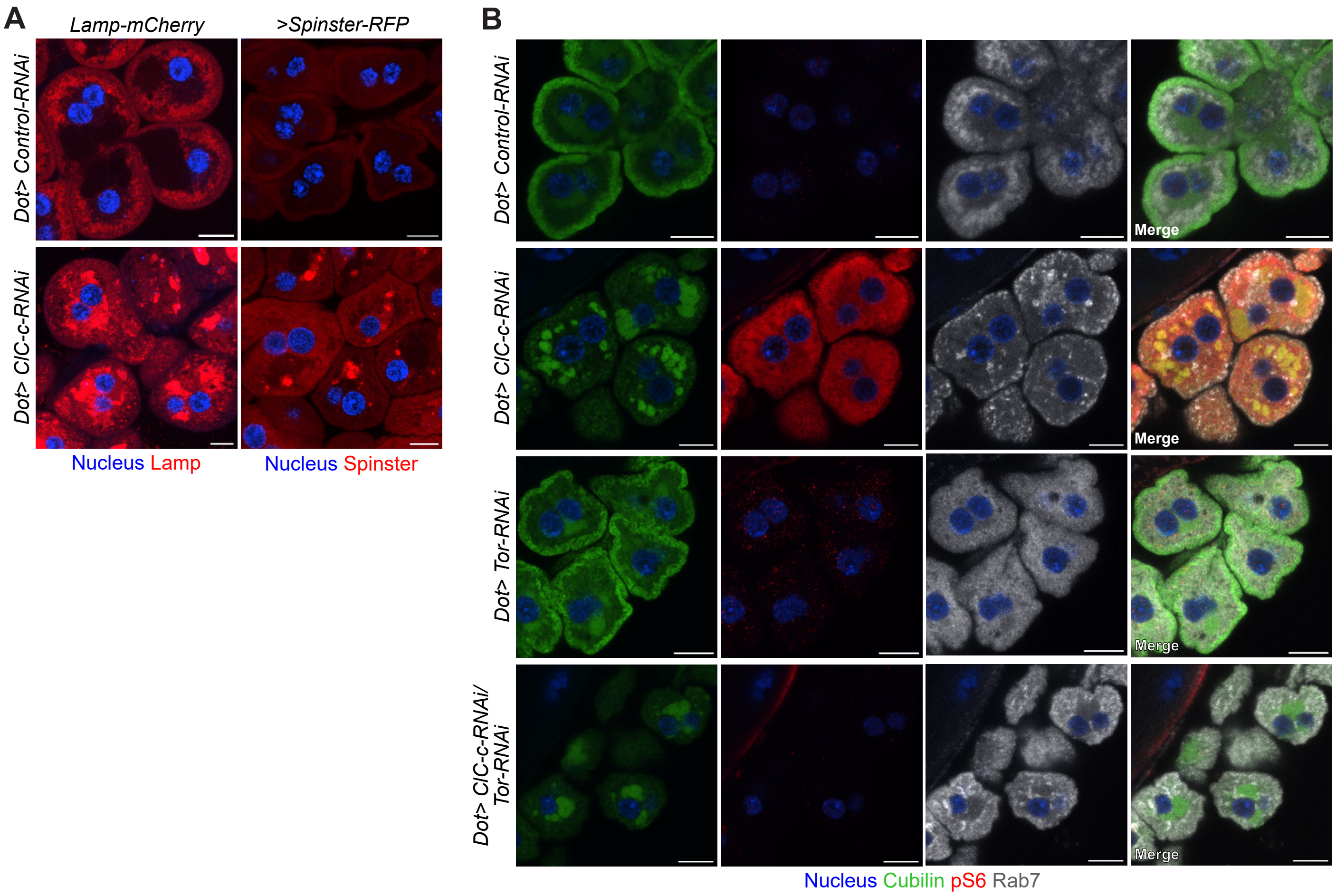


**Suppl. Figure 5: Perinuclear lysosomal clustering in ClC-c KD nephrocytes is independent of Tor signaling.** (A) Visualization of lysosomes through endogenous expression of mCherry-tagged LAMP and overexpression of RFP-tagged Spinster confirms lysosomal identity of perinuclear clusters in *ClC-c*-KD nephrocytes. (B) Nephrocytes co-expressing *ClC-c* and *Tor-RNAi* show that *Tor*-silencing reduces pS6 levels but does not alleviate perinuclear Rab7 clustering or ER retention of Cubilin. Scale bar = 10μm throughout figure.

**Suppl. Fig 6:**

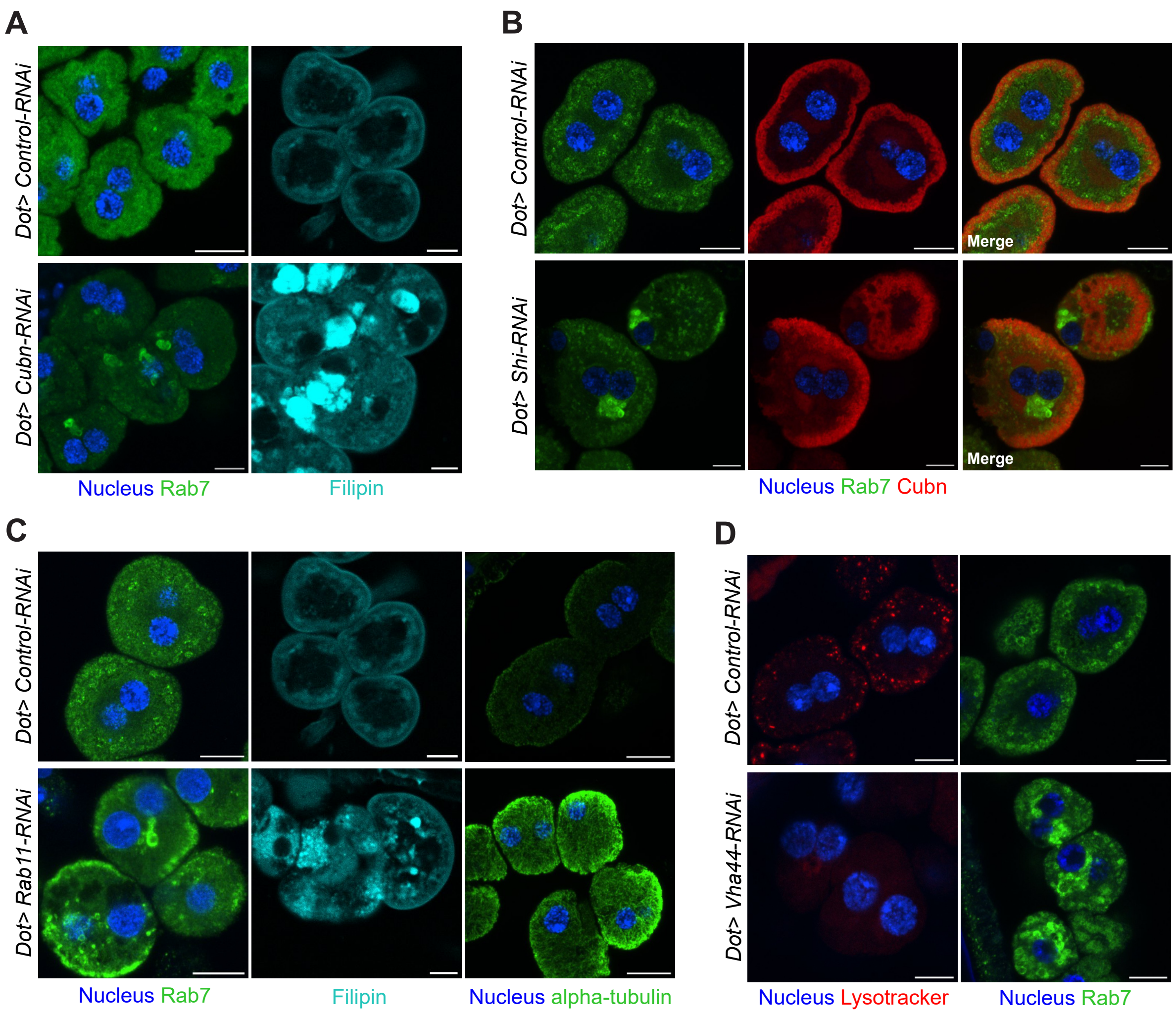


**Suppl. Figure 6: Perinuclear clustering of autolysosomes in *Shi, Cubn, Rab11* and *Vha44* KD.** Ablation of endocytic function through KD of (A) *Cubn* and (B) *Shi* (Dynamin in mammals) leads to formation of filipin-positive perinuclear Rab7-positive clusters, but Cubilin retains its cortical localization upon KD of *Shi*. (C) KD of *Rab11* causes similar perinuclear clustering of cholesterol-rich lysosomes as well as cortical accumulation of microtubules (D). Impaired acidification in *Vha44* KD nephrocytes confirmed through loss of Lysotracker-Red signal. Perinuclear clustering of Rab7 positive vesicles can also be observed upon *Vha44* silencing. Images acquired by confocal microscopy. Scale bar = 10µm throughout figure.
